# Disruption of mitochondrial folate metabolism leads to mitochondrial DNA leakage and activation of apoptotic or inflammatory pathways

**DOI:** 10.1101/2024.07.30.605854

**Authors:** Sinwoo Hwang, Cameron Baker, Martha S. Field

## Abstract

Folate-mediated one-carbon metabolism (FOCM) is required for the biosynthesis of purines, thymidylate (dTMP), and methionine. Maintenance of adequate cellular dTMP levels is essential to preserve the integrity of the nuclear and mitochondrial genomes. Inadequate dTMP production leads to uracil misincorporation into mitochondrial DNA (mtDNA) and impaired mitochondrial function. However, the mechanisms whereby uracil in mtDNA impairs mitochondrial function are uncharacterized. The release of mtDNA into the cytosol acts as a damage-associated molecular pattern (DAMP), resulting in inflammation and cell death. Previously, we reported that disrupting mitochondrial dTMP synthesis through serine hydroxymethyltransferase 2 (*Shmt2*) heterozygosity increases uracil in mtDNA and impairs mitochondrial function in mice. This study aimed to investigate whether impairment of mitochondrial FOCM through *Shmt2* disruption leads to the release of mtDNA into the cytosol. *Shmt2*^+/-^ MEF cells showed a > 2-fold increase in cytosolic mtDNA leakage compared to *Shmt2*^+/+^ cells (p<0.05). There was no significant difference in total mtDNA content between *Shmt2*^*+/+*^ and *Shmt2*^*+/-*^ MEF cells. MEFs with decreased *Shmt2* expression activated apoptosis by the ratio of cleaved caspase-3 to caspase-3. In addition, SHMT2 knock-out (SHMT2 KO) haploid chronic myeloid leukemia (HAP1) cells also exhibited increased cytosolic mtDNA content compared to the wild-type (WT) HAP1 cells. HAP1 lacking *Shmt2* expression activated the cGAS/STING pathway, but suppressed apoptosis, compared to WT HAP1 cells. This study demonstrates that decreased *Shmt2* expression leads to cytosolic mtDNA leakage and that the downstream effects of mtDNA leakage vary by cell type.

## Introduction

Folate, also known as vitamin B9, is an essential cofactor required for folate-mediated one-carbon metabolism (FOCM) [1]. This metabolic pathway provides one-carbon units for biological processes, including biosynthesis of purines, thymidylate (dTMP), and methionine [2]. Disruptions of FOCM are associated with an increased risk of various pathologies such as anemia, cancer, neurodegeneration, and neural tube defects. FOCM is compartmentalized within the cell, taking place in the cytosol, nucleus, and mitochondria [3, 4]. FOCM is essential to maintain an adequate cellular dTMP level to preserve the integrity of the nuclear and mitochondrial genomes. Impaired folate-dependent *de novo* synthesis of dTMP leads to uracil misincorporation into nuclear DNA (nDNA) and mitochondrial DNA (mtDNA), in turn, causing genome instability [5]. In HeLa cells and in mouse liver, folate deficiency elevates uracil in in mtDNA without increasing uracil in nDNA, indicating that mtDNA is more susceptible to disruption of FOCM than is nDNA [6, 7].

Serine hydroxymethyltransferase 2 (SHMT2) facilitates the reversible conversion of THF and serine into 5,10-methylene-THF and glycine within the mitochondria [8]. In addition to SHMT2, mitochondrial biosynthesis of dTMP also requires folate-dependent enzymes synthase (TYMS) and dihydrofolate reductase (DHFR). These enzymes are responsible for converting deoxyuridine monophosphate (dUMP) into dTMP [4, 9, 10]. Therefore, disruption of *Shmt2* expression impairs mitochondrial dTMP synthesis. In mouse liver and cultured cells, *Shmt2* disruption leads to uracil misincorporation in mtDNA and impaired oxidative phosphorylation [11]. The mechanisms whereby uracil misincorporation in mtDNA impairs mitochondrial function have not been elucidated.

Repair of uracil misincorporation by base-excision repair mechanisms can damage DNA by generating strand-breaks and abasic sites during the repair process. MtDNA damage can lead to mtDNA leakage into the cytosol, where mtDNA then acts as a mitochondrial-derived damage-associated molecular pattern (DAMP). Increasing evidence suggests that mitochondria can initiate the innate immune response and apoptosis by releasing mitochondrial DAMPs under various stress conditions [12, 13]. In addition to its role in innate immunity [14], mitochondrial DAMPs are involved in apoptotic cell death through the activation of caspases [15]. Still, it is unclear how mitochondrial DAMPs regulate the balance between immune response and apoptosis. Given the role of SHMT2 in maintenance of mtDNA integrity, the aim of this study was to investigate whether decreased *Shmt2* expression induces mtDNA release into the cytosol and how cytosolic mtDNA release affects downstream processes including apoptosis and the immune response.

## Materials and Methods

### Mouse models, cell lines, and cell culture conditions

*Shmt2*^*+/-*^ whole-body knockout mice (homozygous knockout is embryonic lethal) on a C57BL/6J background were generated as previously described [11]. Mouse embryonic fibroblasts (MEFs) were isolated from C57BL/6J female mice bred to *Shmt2*^*+/-*^ male mice as previously described [11]. HAP1 cells (wild-type) and SHMT2 knock-out HAP1 (*SHMT2* KO) HAP1 cells were obtained from Horizon Discovery. MEF cells were passaged in alpha-minimal essential medium (alpha-MEM; Hyclone Laboratories, Logan, UT, USA) supplemented with 10% FBS (VWR, Radnor, PA, USA) and 1% penicillin/streptomycin (Gibco, Grand Island, NY, USA). HAP1 cells were passaged in Iscove’s Modification of DMEM (IMDM, Corning, NY, USA) supplemented with 10% FBS (VWR, Radnor, PA, USA) and 1% penicillin/streptomycin (Gibco, Grand Island, NY, USA). Cells were cultured at 37 °C in a humidified atmosphere of 5% CO_2_. To identify the effect of folate availability, HAP1 cells were cultured in the folate deficient (0 nM) and the folate sufficient (25 nM) modified DMEM (Hyclone Laboratories, Logan, UT, USA) as previously described (35).

### Preparation of cell extracts

When cultured MEF and HAP1 cells reach 80% confluency in a 6-well plate, the cells were washed with ice-cold phosphate buffered saline (PBS) three times. The cells were collected in 1X radioimmunoprecipitation assay buffer (RIPA buffer; Cell Signaling, Danvers, MA, USA), supplemented with 1 mM PMSF (Cell Signaling, Dancers, MA, USA), 0.5 M EDTA Solution (Thermo Fisher Scientific, Waltham, MA, USA), and Halt Phosphatase Inhibitor Cocktail (Thermo Fisher Scientific, Waltham, MA, USA). The cells were incubated on ice for 20 minutes and centrifuged at 13,000 × g for 10 minutes at 4 °C. The supernatant was transferred to a new 1.5 mL tube. Total protein was quantified by the Pierce BCA protein assay (Thermo Fisher Scientific, Waltham, MA, USA) following manufacturer’s instruction.

### Immunoblotting

Protein lysates were denatured by heating with 4X Laemelli buffer (Bio-Rad, Hercules, CA, USA) for 5 minutes at 95 °C. Samples were loaded onto 4-12% Sodium dodecyl-sulfate polyacrylamide gel electrophoresis (SDS-PAGE) gels for 60-90 minutes in SDS-PAGE running buffer and transferred to Polyvinylidene difluoride (PVDF) membranes (Bio-Rad, Hercules, CA, USA). Membranes were stained with Ponceau S to verify the transfer of proteins. Membranes were blocked using 5% (w/v) non-fat dry milk in 1X Tris-buffered saline and tween-20 (TBS-T) for 1 hour at room temperature. The blocked membranes were incubated in the respective primary antibody at 4 °C overnight. Primary antibodies that were used are as follows: CS (1:1000, ab96600, Abcam, Cambridge, UK), cGAS (1:1000, 31689, Cell Signaling, Danvers, MA, USA), STING (1:1000, 50494, Cell Signaling, Danvers, MA, USA), TBK1 (1:1000, 3504, Cell Signaling, Danvers, MA, USA), Phospho-TBK1 (1:1000, 5483, Cell Signaling, Danvers, MA, USA), Caspase-3 (1:1000, 9662, Cell Signaling, Danvers, MA, USA), Cleaved caspase-3 (1:1000, 9664, Cell Signaling, Danvers, MA, USA), and GAPDH (1:2118S, Cell Signaling, Danvers, MA, USA). Membranes were washed 3 times with 1X TBS-T for 5 minutes each and incubated in the appropriate horseradish peroxidase-conjugated secondary antibody for 2 hours at room temperature, followed by washing 3 times with 1X TBS-T. To detect antibody, an enhanced chemiluminescent reagent (SuperSignal west femto maximum sensitivity substrate, Thermo Fisher Scientific, Waltham, MA, USA) was used at a 1:10000 dilution. Blots were imaged using FluorChem E imaging system (Protein Simple, San Jose, CA, USA). Densitometry was performed with Fiji software using GAPDH as the control.

### Sub-fractionation of cellular compartments (Mitochondria)

Cultured MEF cells (5 × 10^6^) or HAP1 (1 × 10^7^) were pelleted by centrifugation at 500 × g for 10 minutes at 4 °C. The pellets were washed with 0.9 % sodium chloride (NaCl) and pelleted again by centrifugation at 500 × g for 10 minutes at 4 °C. Qproteome mitochondria isolation assay (Qiagen, Hilden, Germany) was performed to extract high-purity mitochondria. The pellets were resuspended in ice-cold Qproteome lysis buffer supplemented with 100X protease inhibitor solution and incubated for 10 minutes at 4 °C on an end-over-end shaker. The lysates were pelleted by centrifugation at 1000 × g for 10 minutes at 4 °C and resuspended in ice-cold Qproteome disruption buffer. The lysates were disrupted with a blunt-ended needle and a syringe and centrifugated at 1000 × g for 10 minutes at 4 °C. The supernatant was transferred to a fresh 1.5 mL tube, followed by another centrifugation at 6000 × g for 10 minutes at 4 °C to obtain mitochondria-containing pellets. To obtain high-purity mitochondria, the pellets were resuspended in Qproteome mitochondrial purification buffer and pipetted on top of layers of purification buffer and disruption buffer, followed by a centrifugation at 14000 × g for 15 minutes at 4 °C. The mitochondrial pellets were washed three times with Qproteome mitochondrial storage buffer and prepared for further experiments.

### Sub-fractionation of cellular compartments (Cytosol)

Cultured cells were washed three times with ice-cold PBS and pelleted by centrifugation at 500 × g for 5 minutes at 4 °C. Cell pellets were resuspended in cytosolic extraction buffer (150 mM NaCl, 50 mM HEPES pH 7.4, 25 μg/ml digitonin) and homogenized using a 2 mL Dounce grinder with a tight-fitting pestle (Kimble Chase, Vineland, NJ, USA) for 10 minutes at 4 °C. Lysates were centrifuged at 10000 × g for 15 minutes at 4 °C. The supernatant was transferred to a fresh 1.5 mL tube, followed by another centrifugation at 12000 × g for 15 minutes at 4 °C. The cytosolic fraction was isolated by collecting supernatant into a fresh 1.5 mL tube. The purity of cytosolic fraction without mitochondrial contamination was determined by immunoblotting analysis with CS (1:1000, ab96600, Abcam, Cambridge, UK)

### Measurement of mitochondrial DNA content

Total genomic DNA was isolated from whole, mitochondrial, and cytosolic cellular fractions respectively, using the QIAamp DNA Micro Kit (Qiagen, Hilden, Germany) following manufacturer’s instructions. DNA purity and concentration were quantified using Nanodrop spectrophotometer (Thermo Fisher Scientific, Waltham, MA, USA). To determine mtDNA copy number, real-time quantitative polymerase chain reaction (qPCR) was performed with the LightCycler 480 SYBR-Green I Master (Roche Diagnostics, Basel, Switzerland) and primers specific to the mitochondrial D-loop region that is not inserted into nuclear DNA region (non-NUMT). Nuclear DNA encoding *Tert* primers were used for normalization. PCR amplification was performed with a 10-minutes denaturation step at 95°C, followed by 40 cycles of 95°C for 10 seconds, 60°C for 30 seconds, and 72°C for 10 seconds. Primer sequences are listed as follows:

Mouse D-loop forward: 5’-AATCTACCATCCTCCGTGAAACC-3’

Mouse D-loop reverse: 5’-TCAGTTTAGCTACCCCCAAGTTTAA-3’

Mouse *Tert* forward: 5’-CTAGCTCATGTGTCAAGACCCTCTT-3’

Mouse *Tert* reverse: 5’-GCCAGCACGTTTCTCTCGTT-3’

Human D-loop forward: 5’-GTTTATGTAGCTTACCTCCTC-3’

Human D-loop reverse: 5’-TTGTTTATGGGGTGATGTGAG-3’

Human *Tert* forward: 5’-TCACGGAGACCACGTTTCAAA-3’

Human *Tert* reverse: 5’-TTCAAGTGCTGTCTGATTCCAAT-3’

### Statistical analysis

Statistical analysis was performed using GraphPad Prism 8.0. The Welch’s t-test was used to determine whether there is a significant difference between the means of *Shmt2*^*+/+*^ and *Shmt2*^*+/-*^ MEFs, and wild-type HAP1 and SHMT2 KO HAP1. A *p*-value of 0.05 or less was considered statistically significant.

## Results

### Reduced *Shmt2* expression leads to mtDNA leakage into the cytosol without changing total mtDNA content

To examine mtDNA leakage into the cytosol, cytosol was isolated from the cultured *Shmt2*^*+/+*^ and *Shmt2*^*+/-*^ MEF cells. Using the primers specific to the mitochondrial D-loop region that is not inserted into nuclear DNA (non-NUMT) and nuclear DNA encoding *Tert* primers as control, cytosolic mtDNA content was measured by qPCR. Citrate synthase, a mitochondrial matrix enzyme, is commonly used as a maker for the presence of intact mitochondria (i.e. lack of mitochondrial contamination in cytosolic fraction). As shown in Figure 1A and 1D, respectively, citrate synthase was only observed in the total cell lysate and not in the cytosol of MEF (*Shmt2*^*+/+*^ and *Shmt2*^*+/-*^) and HAP1 (wild-type and *SHMT2* KO) cells, confirming the purity of the cytosolic fractions. There was no significant difference in total cellular mtDNA content between *Shmt2*^*+/+*^ and *Shmt2*^*+/-*^ MEF cells (Figure 1B), consistent with previous results [11]. However, the *Shmt2*^*+/-*^ MEF cells showed a > 2-fold increase in cytosolic mtDNA leakage compared to the *Shmt2*^*+/+*^ group (Figure 1C). Similarly, *SHMT2* KO HAP1 cells exhibited higher levels of mtDNA content in the cytosol compared to the wild-type HAP1 cells (Figure 1F). As observed in MEF cells (Figure 1B), there was no significant difference in total mtDNA between wild-type and *SHMT2* KO HAP1 cells (Figure 1E). In addition, wild-type HAP1 cells were cultured in the folate-deficient modified cell culture medium to determine if the folate availability affects the cytosolic mtDNA leakage. No significant change was observed in the cytosolic mtDNA content between the control and folate-deficient media groups (Figure 1G), suggesting that exposure to folate-deficient culture medium does not increase mtDNA leakage to the cytosol.

**Figure 1.**
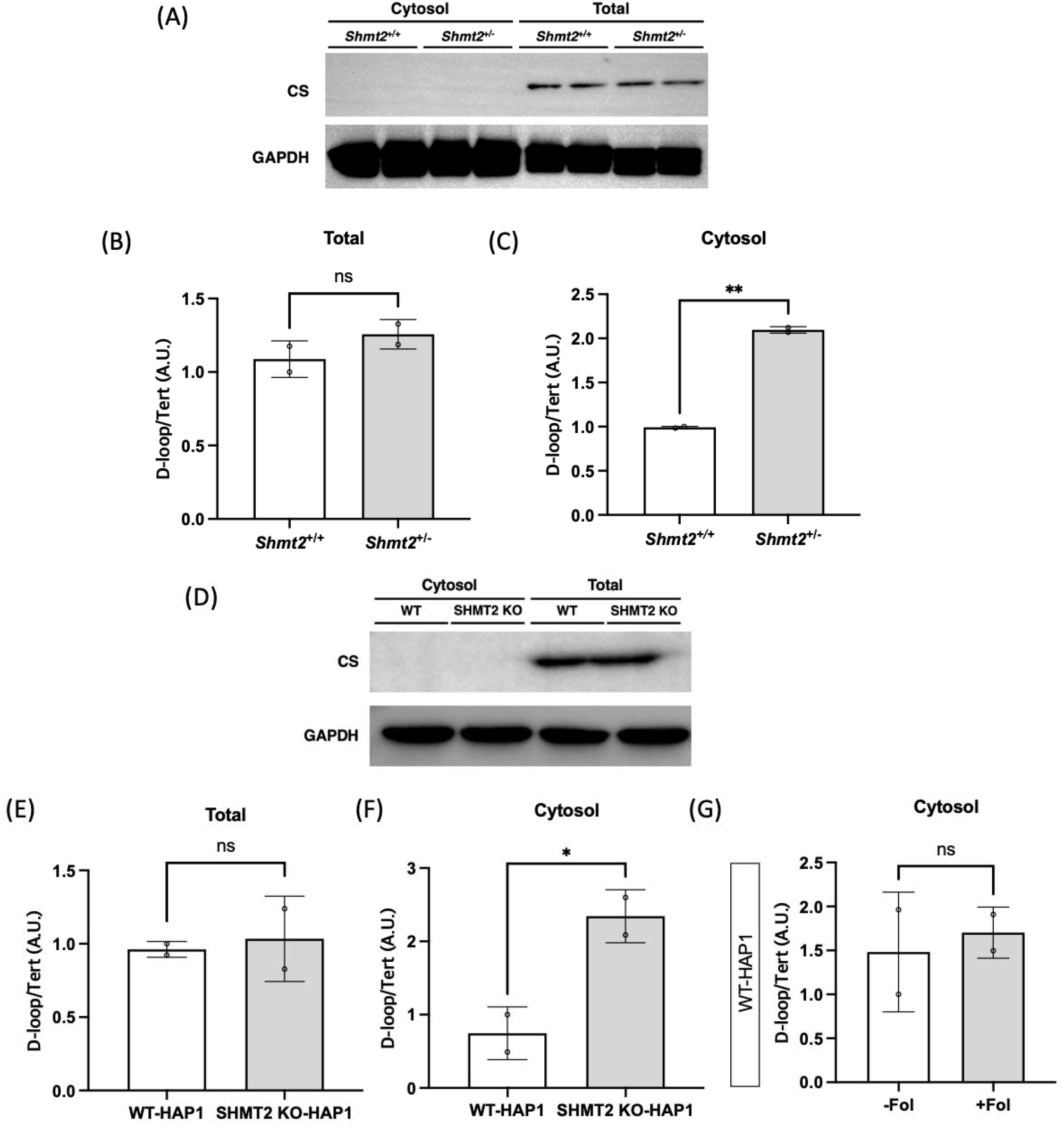
Decreased *Shmt2* expression leads to mitochondrial DNA leakage into the cytosol without changing cellular mtDNA content in MEFs and HAP1 cells. (A) Representative western blots of CS protein expression in subcellular fractionation of cytosol and whole cell lysates. (B) Total mtDNA content and (C) cytosolic mtDNA content in *Shmt2* MEF cells. (D) Representative western blots of CS protein expression in subcellular fractionation of cytosol and whole cell lysates. (E) Total mtDNA content and (F) cytosolic mtDNA content of Shmt2 wild-type and knock-out HAP1 cells cultured in IMDM. (G) Cytosolic mtDNA content of wild-type HAP1 cells in folate modified media. Densitometry was performed with Fiji software using GAPDH as a control. A Welch’s t test was performed to determine the statistical significance. AU, arbitrary units; CS, citrate synthase; D-loop, displacement loop; GAPDH, glyceraldehyde-3-phosphate dehydrogenase; Tert, telomerase reverse transcriptase. ns denotes statistically not significant, * denotes P < 0.05, and ** denotes P < 0.01.

### *Shmt2* heterozygosity-induced mtDNA leakage increases apoptosis in MEFs but suppresses apoptosis in HAP1 cells

Heterozygous disruption of *Shmt2* expression in MEF cells lead to an increase in cytosolic mtDNA leakage (Figure 1). Since cytosolic mtDNA leakage is linked to both and inflammation, we investigated whether reduced expression of *Shmt2* increases pro-apoptotic activity by measuring the ratio of cleaved caspase-3 to caspase-3 in whole-cell lysates of *Shmt2*^*+/+*^ and *Shmt2*^*+/-*^ MEF cells. *Shmt2*^+/-^ MEF cells had a significantly higher level of the cleaved caspase-3 compared to *Shmt2*^*+/+*^ MEF cells. Furthermore, the ratio of cleaved caspase-3 to pro-caspase-3 was more than 10-fold higher in *Shmt2*^*+/-*^ MEF cells compared to *Shmt2*^*+/+*^ MEF cells (Figure 2A-C). In contrast, *SHMT2* KO-induced cytosolic mtDNA in HAP1 cells was associated with decreased apoptotic pathway activation compared to WT HAP1 cells (Figure 2D-F). Taken together, these results suggest that *Shmt2* heterozygosity-induced mtDNA leakage into the cytosol is associated with apoptosis in MEF, but not HAP1 cells.

**Figure 2.**
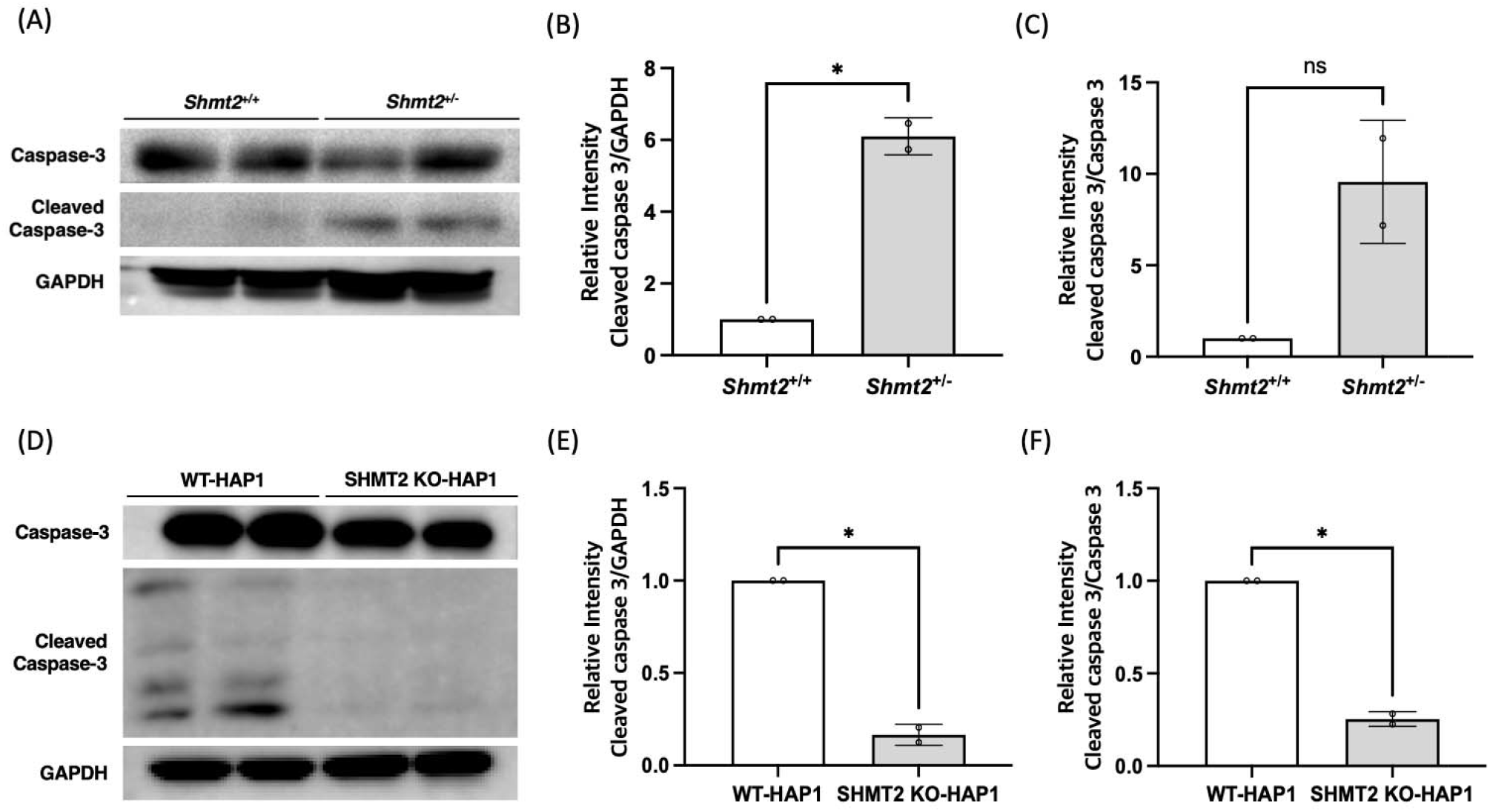
*Shmt2* heterozygosity-induced mtDNA leakage increases apoptosis in MEF cells but suppresses apoptosis in SHMT2 KO HAP1 cells. (A) Representative western blot of caspase-3 and cleaved caspase-3 and quantification graphs of (B) cleaved caspase-3 and (C) the ratio of cleaved caspase-3 to caspase-3 in *Shmt2*^*+/+*^ and *Shmt2*^*+/-*^ MEF cells. (D) Representative western blot of caspase-3 and cleaved caspase-3 and quantification graphs of (E) cleaved caspase-3 and (F) the ratio of cleaved caspase-3 to caspase-3 in WT or SHMT2 KO HAP1 cells. Densitometry was performed with Fiji software using GAPDH as a control. A Welch’s t test was performed to determine the statistical significance. ns denotes statistically not significant and * denotes P < 0.05.

### Cytosolic mtDNA is associated with activation of the cGAS/STING pathway in HAP1 cells lacking *SHMT2* but not with *Shmt2* heterozygosity in MEF cells

Previous studies have demonstrated that apoptotic caspases attenuate cGAS/STING-mediated inflammatory signals triggered by mtDNA [16]. In light of our discovery that the release of mtDNA into the cytosol due to *Shmt2* heterozygosity is linked to apoptosis in MEF cells (Figure 2), we hypothesized that the activity of the cGAS/STING pathway would be decreased in *Shmt2*^*+/-*^ MEF cells. The protein levels of cGAS and STING were evaluated in *Shmt2*^*+/+*^ and *Shmt2*^*+/-*^ MEF cells using western blotting. No notable differences were observed between two groups (Figure 3A-C). Interestingly, SHMT2 KO HAP1 cells exhibited significant higher STING expression and an increase in the ratio of pTBK-1/TBK-1, compared to the wild-type HAP1 cells (Figure 3D-3F). This suggests that the *Shmt2* heterozygosity-induced mtDNA release into the cytosol does not activate cGAS/STING pathway in MEF cells; conversely, HAP1 cells lacking SHMT2 expression was associated with cGAS/STING activation.

**Figure 3.**
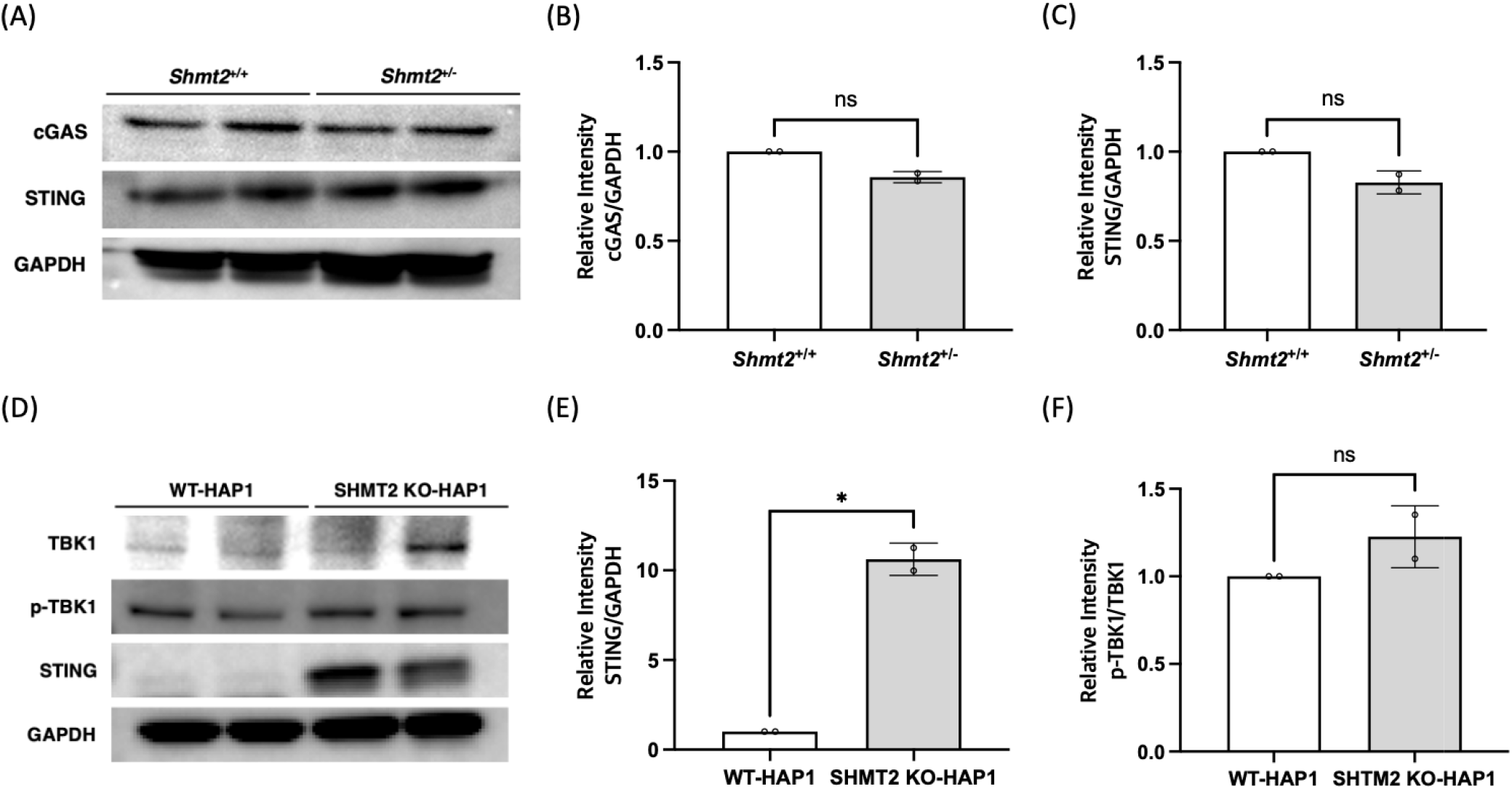
*Shmt2* loss activates cGAS/STING-mediated inflammation in *SHMT2* KO HAP1 cells. (A) Representative western blot of cGAS and STING and the quantification graphs of (B) cGAS and (C) STING in *Shmt2*^*+/+*^ and *Shmt2*^*+/-*^ MEF cells. As the samples are the same as those shown in Figure 2A, the GAPDH western blot image was reused in Figure 3A as a control. (D) Representative western blot of TBK1, p-TBK1, and STING and the quantification graphs of (E) STING and (F) the ratio of p-TBK1/TBK1 in wild-type and SHMT2 KO HAP1 cells. As the samples are the same as those shown in Figure 2D, the GAPDH western blot image was reused in Figure 3D as a control. Densitometry was performed with Fiji software using GAPDH as a control. A Welch’s t test was performed to determine the statistical significance. cGAS, cyclic GMP-AMP synthase; STING, stimulator of interferon gene; TBK1, tank binding kinase 1; p-TBK1, phosphorylated tank binding kinase 1. ns denotes statistically not significant and * denotes P < 0.05.

### Conclusions and discussion

Mitochondria are essential organelles within eukaryotic cells that play a crucial role in energy production and cellular respiration [17-19]. SHMT2 is a folate-dependent enzyme that is integral to mitochondrial FOCM. Mitochondrial FOCM is essential for various cellular processes, including the biosynthesis of formate for cytosolic/nuclear FOCM, mitochondrial dTMP production, and intermediates required for mitochondrial protein translation [4]. Recent studies have identified biallelic human *SHMT2* variants that result in decreased in SHMT2 expression in human fibroblasts with age [20, 21]. This has led to a growing interest in understanding the cellular implications of perturbed mitochondrial FOCM.

Moreover, research on the *Shmt2* gene and its association with mtDNA stability and mitochondrial function has attracted significant attention. Previously, we demonstrated that *Shmt2* heterozygosity results in uracil misincorporation into mtDNA and impaired mitochondrial function without affecting the total mtDNA mass [11]. However, the impact of *Shmt2* heterozygosity on other aspects of mitochondrial biology remains unclear. In this study, we aimed to investigate whether *Shmt2* heterozygosity could lead to mtDNA leakage, activation of apoptosis, increased mtDNA common deletions, and accumulated deleted mtDNA within the mitochondrial compartment.

Here, we demonstrated that reduced expression of *Shmt2* led to mtDNA leakage into the cytosol in MEF cells. Interestingly, there was no significant change in total mtDNA content comparing wild-type and *Shmt2*^*+/-*^ MEF cells (Figure 1C), suggesting that impaired mitochondrial FOCM induced by decreased *Shmt2* may lead to leakage of mtDNA into the cytosol by altering mitochondrial membrane integrity. These findings corroborate the previous research of Fiddler *et al*. (2021), demonstrating that *Shmt2* heterozygous MEF cells cultured in low-folate media had decreased mitochondrial membrane potential [11].

As mentioned above, reduced *Shmt2* expression leads to mitochondrial FOCM disruption, resulting in the accumulation of uracil misincorporation in mtDNA of mouse liver [11]. Repairing uracil in nuclear DNA is known to induce genome instability through single- and double-strand breaks [22]. However, the consequences of uracil misincorporation into mtDNA remain uncharacterized. Nonetheless, mtDNA breaks – expected during uracil excision repair in mitochondria – are known to cause mtDNA deletions [23]. The findings presented here suggest that uracil-induced mtDNA damage may lead to mtDNA release into the cytosol.

MtDNA is increasingly recognized as a regulator of immune responses and apoptotic cell death [14, 24]. Mitochondrial DAMPs are endogenous molecules that can alert the immune system to potential threats, such as tissue damage or infection [25]. In particular, mtDNA leakage is a crucial component of this alarm system, triggering the activation of PRRs to initiate a cascade of immunological events to repair or eliminate threats to the host organism [14]. Our hypothesis posited that decreased *Shmt2* expression would lead to mtDNA leakage into the cytosol and trigger an innate immune response. Contrary to this initial hypothesis, no significant difference was observed in the expression levels of cGAS/STING between wild-type and *Shmt2*^*+/-*^ MEF cells (Figure 3). The cGAS/STING signaling pathway is one type of the innate immune system that detects the presence of cytosolic DNA. Specifically, cGAS recognizes and binds to double-stranded DNA (dsDNA), catalyzing the synthesis of cyclic guanosine monophosphate-adenosine monophosphate (cGAMP) from ATP and GTP [26, 27]. The cGAMP, in turn, activates the STING protein as a second messenger [28]. Subsequently, STING activates TANK-binding kinase 1 (TBK1) through its C-terminal PLPLRT/SD motif [29]. TBK1 phosphorylates the interferon regulatory factor 3 (IRF3) [30, 31]. Dimerized IRF3 translocate into the nucleus and induces the expression of type I IFNs [30].

It may be possible that *Shmt2* heterozygosity-induced mtDNA leakage is not sufficient to activate the cGAS/STING pathway. Alternatively, there may be other cellular mechanisms that can compensate for the increased mtDNA leakage in *Shmt2* heterozygous MEF cells and maintain a balanced immune response to prevent cGAS/STING activation. Interestingly, we found that *Shmt2* heterozygosity-induced mtDNA leakage can promote apoptosis through a cascade of caspase, instead of activating cGAS/STING-mediated inflammation (Figure 2). This finding is consistent with previous research that apoptotic caspases suppress type I IFN production via the cleavage of cGAS [32]. In contrast, *SHMT2* KO HAP1 cells also exhibit increased mtDNA leakage to the cytosol, but in this cell type the cGAS/STING pathway is activated as a result of *SHMT2* loss (Figure 2).

The findings presented in this study offer novel insights into the mechanisms underlying SHMT2 disruption-induced impairments in FOCM, which can lead to mitochondrial dysfunction and immune response activation and/or cell death. It is important to note that mtDNA release as a result of impaired mitochondrial FOCM is observed only in response to reduced *SHMT2* expression, and not as a result of exposure to folate-deficient culture medium. Both folate deficiency and *Shmt2* disruption were shown to increase uracil accumulation in mtDNA and impair mitochondrial function in previous studies [11, 33], suggesting that decreased SHMT2 may cause cytosolic mtDNA leakage through mechanisms other than just impairing FOCM such as impaired mitochondrial translation or impaired redox balance [4]. *SHMT2* expression decreases with aging in human tissues [21], highlighting the need to better understand the mechanisms linking decreased *SHMT2* expression, mtDNA leakage, and immune activation in the context of aging. Future studies should focus on determining whether cytosolic mtDNA can initiate the innate immune response through TLR9 or inflammasome-mediated pathways in addition to the cGAS/STING pathway. In addition, it remains unclear which membrane pore is responsible for the mtDNA release under as a result of reduced *Shmt2* expression. Therefore, future work should focus on investigating the role of SHMT2 in cell-free mtDNA-mediated cell death across a range of tissues and determine the molecular mechanisms that cause mtDNA leakage.

## Acknowledgements

The authors thank Luisa Castillo and Katarina Heyden for technical assistance.

## Author contributions

SH and MSF designed research. SH and CB conducted research. SH, CB and MSF analysed data. SH prepared the figures. SH and MSF prepared the manuscript. MSF has primary responsibility for the final content. All authors reviewed and approved the final manuscript.

